# Perturbations in dynamical models of whole-brain activity dissociate between the level and stability of consciousness

**DOI:** 10.1101/2020.07.02.185157

**Authors:** Yonatan Sanz Perl, Carla Pallavicini, Ignacio Pérez Ipiña, Athena Demertzi, Vincent Bonhomme, Charlotte Martial, Rajanikant Panda, Jitka Annen, Agustín Ibañez, Morten Kringelbach, Gustavo Deco, Helmut Laufs, Jacobo Sitt, Steven Laureys, Enzo Tagliazucchi

## Abstract

Consciousness transiently fades away during deep sleep, more stably under anesthesia, and sometimes permanently due to brain injury. The development of an index to quantify the level of consciousness across these different states is regarded as a key problem both in basic and clinical neuroscience. We argue that this problem is ill-defined since such an index would not exhaust all the relevant information about a given state of consciousness. While the level of consciousness can be taken to describe the actual brain state, a complete characterization should also include its potential behavior against external perturbations. We developed and analyzed whole-brain computational models to show that the stability of conscious states provides information complementary to their similarity to conscious wakefulness. Our work leads to a novel methodological framework to sort out different brain states by their stability and reversibility, and illustrates its usefulness to dissociate between physiological (sleep), pathological (brain-injured patients), and pharmacologically-induced (anesthesia) loss of consciousness.

## Introduction

Human consciousness can be understood in terms of its contents, but also as a state extended in time. The contents of consciousness are frequently investigated from a functional perspective, combining task-based paradigms from cognitive neuroscience with different neuroimaging methods to reveal the brain mechanisms associated with explicit or implicit reports of conscious awareness (1). The study of temporally extended conscious states is more elusive, with different authors agreeing more often on specific examples than on broad and clear definitions (2–5). Some examples include the different stages of the human wake-sleep cycle (6), the acute effects of anesthesia (7), and post-comatose disorders of consciousness (8). These states cannot be defined by the specific nature of their content; instead, our intuition suggests that they involve overall reductions in the intensity or level of consciousness, perhaps up to the point of becoming completely void of subjective experiences. A natural question emerges from these considerations: how can the level of consciousness be estimated from third-person measurements, and what is the validity of this estimation?

Several unidimensional scales have been developed and implemented for the purpose of measuring the level of consciousness. Some are based on the observation and quantification of behaviour, such as the scales used during surgery to determine the depth of anesthesia, or those employed by neurologists to diagnose the severity of disorders of consciousness in brain-injured patients (9). Others are obtained from recordings of spontaneous or evoked brain activity, either by following a data-driven approach or by computations informed by theoretical developments (10–15). For instance, information integration theory (IIT) posits that the level of consciousness corresponds to the amount of integrated information (Φ) in the brain (16), which can be approximated by different metrics of complexity and data compressibility (17). These and other examples present the common feature of collapsing whole-brain activity data into a single numerical index that is expected to correlate with the outcome of behavioral scales, akin to a “thermometer” capable of objectively measuring the level of consciousness taking a state of healthy wakefulness as a reference point (18).

Leaving issues of practicality aside, it seems questionable that a single number can adequately describe the global state of human consciousness. The brain is a highly complex system composed of 10^10^ nonlinear units (neurons) interacting in 10^15^ sites (synapses) (19). Considering this astonishing level of complexity, it is surprising that the brain self-organizes into a discrete and reduced number of behaviorally distinct states (20), let alone that these states can be placed along a unidimensional continuum parametrized by the level of consciousness (5). In reality, most numerical constructs associated with human brain and behaviour present multiple independent factors - consider, for example, the leading models of personality, intelligence, and mental disorders, all of which are multidimensional (21–23). Analogously, states of consciousness can be described by factors related to content (e.g. the level of sensory gating) and function (e.g. the degree of global availability of information for cognitive processing), a view that can be extended to states observed during certain psychiatric conditions, or arising as a consequence of pharmacological stimulation (24). In turn, each of these dimensions implicates specific brain functional systems, as opposed to the global and emergent character of metrics related to complexity and information integration. Still, numerical metrics for consciousness could be valid as first approximations useful to assist clinicians in the diagnosis and prognosis of difficult cases (8).

In contrast to this view, we propose that numerical metrics are essentially insufficient - even when considered as approximations. A way of framing this insufficiency is noting that consciousness can be characterized by descriptive and perturbational dimensions or, equivalently, by the actual and potential state of the brain. The first dimension is related to the question *“how does this state feel like and what sort of behavior does it entail?”*, while the second addresses the stability of the state, thus answering the question: *“how will this state behave if perturbed?”*. The perturbational dimension is uncorrelated with complexity-based metrics, since both deep sleep and general anesthesia present marked reductions in complexity and information integration (15,25), yet the first can spontaneously revert to wakefulness upon sensory stimulation, while the second is associated with a more persistent state of unresponsiveness. An analogy with the mechanical systems studied in physics can be useful to consolidate the difference between both ways of describing a global brain state. Following this analogy, states of consciousness are identified with equilibrium points, and the level of consciousness corresponds to the mechanical energy associated with those points (26). As in the physical analogy, however, the level of consciousness fails to completely characterize present and future behavior. For this purpose, the stability must also be taken into account; for instance, by investigating how the system reacts to different external perturbations.

We pursued this line of inquiry by investigating functional magnetic resonance imaging (fMRI) data corresponding to three different states of reduced consciousness: sleep, propofol anesthesia, and post-comatose disorders of consciousness in brain injured patients. First, we compared the similarity between these states using machine learning classifiers, thus constructing metrics that reflect the level of conscious awareness relative to reference states of healthy wakefulness. Next, we assessed whether transitions between these states could be induced by external stimulation in whole-brain dynamical system models of fMRI data (27), leading to a perturbation similarity metrics.

## Results

### Analysis overview

We explored the similarities and differences between brain states associated with varying levels of consciousness, including wakefulness (W), three progressively deeper sleep stages (N1, N2, N3), propofol-induced sedation (S) and anesthesia (LoC), and in patients suffering from disorders of consciousness (DoC), all diagnosed as unresponsive wakefulness syndrome (UWS) or in the minimally conscious state (MCS). These states were compared using four different distance metrics. The “classification distance” between two states was obtained by training a random forest classifier to distinguish the first state from wakefulness, and assessing its generalization accuracy (i.e. transfer learning) when distinguishing the second state from wakefulness, using whole-brain functional connectivity (FC) between blood-oxygen-level-dependent (BOLD) signals as input (Pallavicini et al., 2019). The “connectivity correlation distance” between two states was based on computing the linear correlation coefficient between regional FC profiles from both states after subtraction of wakefulness FC as a baseline. A high correlation implied that the FC profile of the region changed similarly between both states relative to wakefulness. This distance was then defined as the proportion of regions in the whole-brain parcellation presenting a significant (p<0.05, Bonferroni corrected) correlation with R ≥ 0.5.

The remaining two definitions were based on the results of a whole-brain computational model of brain activity, constructed by coupling regional dynamics with structural connectivity estimated from DTI data (Fig. 1) (28). The dynamics of each region were given by the normal mode of a Hopf bifurcation (also known as a Stuart-Landau nonlinear oscillator), presenting three qualitatively different regimes depending on a single bifurcation parameter: steady dynamics governed by noise (**a**<0), self-sustained oscillations at the fundamental frequency of the regional empirical BOLD time series (**a**>0), and unstable behavior switching back and forth between these two regimes (**a**≈0) (29). Local bifurcation parameters were optimised to reproduce the empirical FC of each state, with the constraint that regions located within different resting state networks (RSNs) (30) contributed as independent anatomical priors to parameter variation (28). Afterwards, the “model parameter distance” between two states was computed as the euclidean distance between the associated set of optimal bifurcation parameters obtained after fitting the model to the empirical FC using genetic algorithms (i.e. one local parameter per region in the whole-brain parcellation). Finally, the “perturbational distance” was determined from the behavior of the whole-brain model against simulated external oscillatory perturbations. This framework consists of fitting the whole-brain model to the empirical FC of each brain state and then applying an *in silico* stimulation protocol to assess the likelihood of inducing transitions between pairs of states (27,31). The procedure followed to construct the model and its sources of empirical information are described in Fig. 1. We then simulated external perturbations using an additive periodic forcing term of variable amplitude incorporated to the dynamical equations of each pair of homotopic regions, and evaluated whether the perturbation increased the similarity between the simulated FC and the empirical FC of another target state (27,28). For example, we evaluated whether external stimulation applied to the model fitted to wakefulness FC could displace the simulated FC towards that of sleep, anesthesia or brain injured patients, and vice-versa. If affirmative, we interpreted that a transition could be induced between both states, leading to a low “perturbational distance” value.

**Figure 1.**
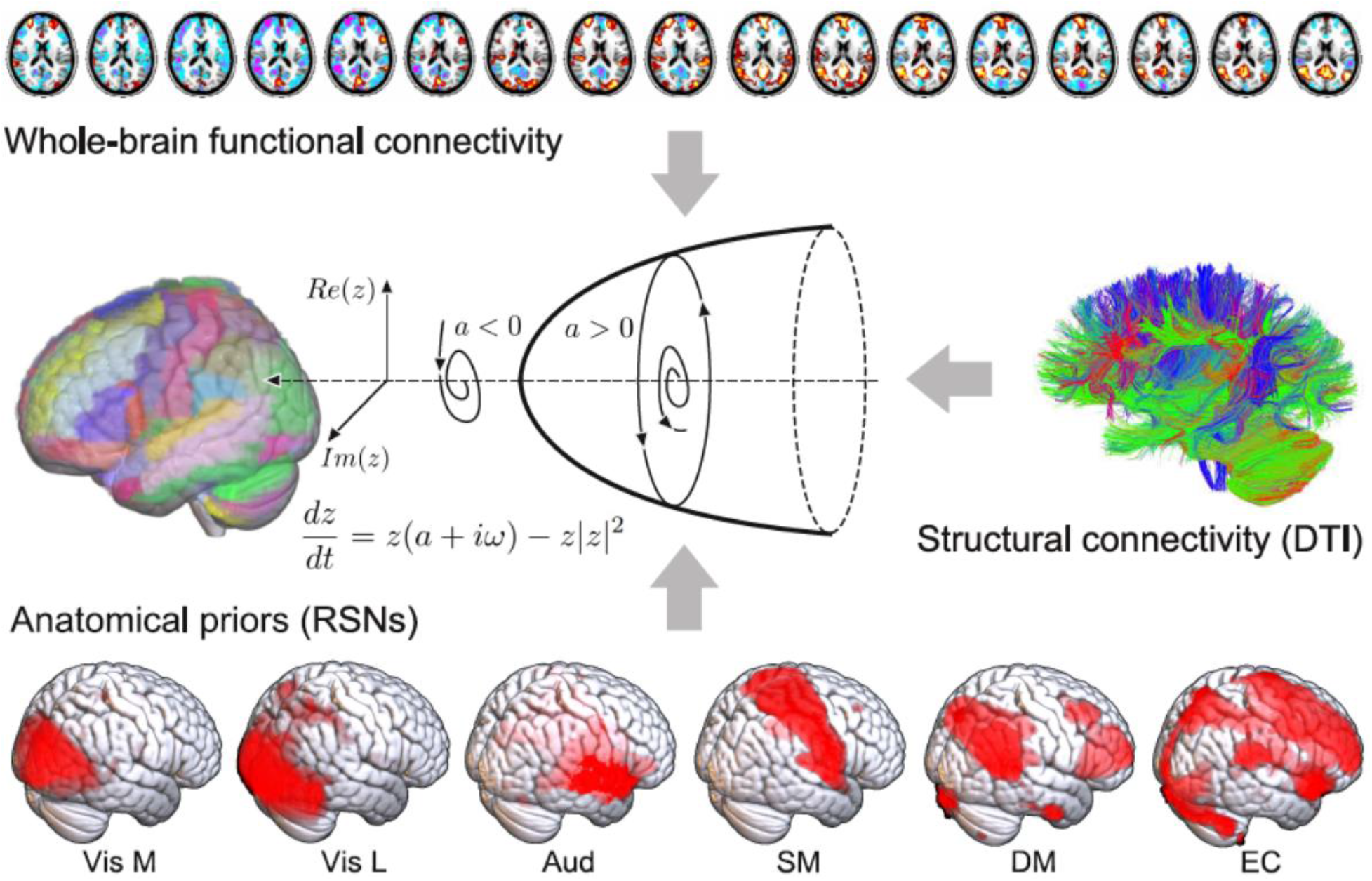
Procedure followed to construct the whole-brain computational model. The dynamics of each node in the structural connectivity matrix are represented by a Hopf bifurcation with three possible dynamical regimes depending on the value of the bifurcation parameter: stable fixed-point (**a**<0), stable limit cycle (**a**>0) and a bifurcation between both regimes (**a**≈0). The local bifurcation parameters are optimized to reproduce the empirical FC matrix computed from fMRI data acquired during different states of consciousness, and constrained in their variation by the six RSNs (anatomical priors) reported in Beckmann et al. (2006): Vis M (medial visual), Vis L (lateral visual), Aud (auditory), SM (sensorimotor), DM (default mode), EC (executive control).

The first three metrics are data-driven and can be computed directly from the empirical fMRI data, or from the inferred model parameters without addition of external stimulation, hence they can be considered descriptive metrics. The fourth metric is perturbational, since it measures whether external stimulation can drive simulated whole-brain FC between patterns typical of different states of consciousness.

### Classification distance between N3, LoC and UWS

We started by studying the similarity between states associated with deepest unconsciousness in our dataset: N3 sleep, LoC and the UWS group of patients. As an exploratory first step, we calculated the average difference in FC for each state vs. wakefulness. These differences are shown in Fig. 2A, both in matrix form and as anatomical renderings of the functional connections associated with the top and bottom 5% differences. A similar pattern of FC changes is evident for N3 and LoC (correlation between FC difference matrices: R=0.65), consisting of reduced FC in occipital and parietal regions, and increased FC in frontal regions. On the other hand, UWS patients did not present such clear patterns, with high magnitude FC differences scattered throughout the whole brain (UWS vs. LoC: R=-0.1; UWS vs. N3: R = −0.1). This suggests that FC changes relative to conscious wakefulness during N3 sleep and LoC present substantial similarities, but are generally different from those seen in UWS patients.

**Figure 2.**
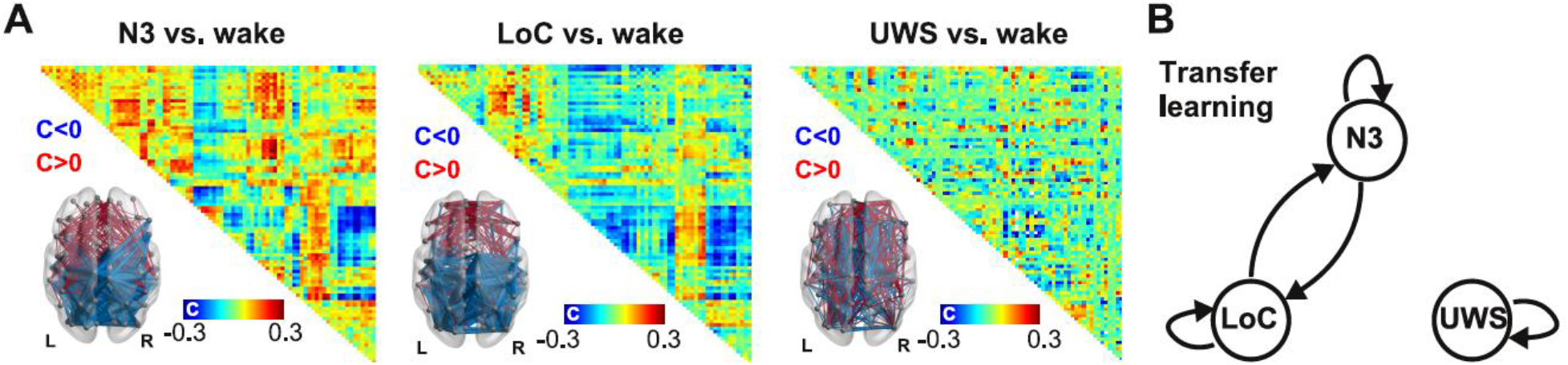
Significant transfer learning accuracy between physiological and pharmacologically-induced states of unconsciousness (N3 sleep and LoC), but not between them and pathological states of unconsciousness (UWS). (A) Average FC differences for N3 vs. wake (left), LoC vs. wake (center), and UWS vs. wake (right), together with anatomical renderings of the top (red) and bottom (blue) 5% functional connections associated with the largest difference between states in absolute value. (B) Nodes in the diagram represent different brain states (N3, LoC, and UWS) and the arrows between them indicate that a machine learning classifier trained to distinguish the source state from W presented significant transfer learning accuracy when distinguishing the target state from W (p<0.05, random label shuffling with 1000 iterations, Bonferroni corrected).

Next, we used individual subject FC data to train and evaluate three random forest classifiers to distinguish W from N3, LoC and UWS. After training and testing by cross-validation, each classifier was applied to recognize the other two brain states from the corresponding W data (i.e. transfer learning was assessed). Arrows in Fig. 2B indicate significant transfer learning classification accuracy; for example, an arrow from LoC to N3 indicates that a random forest classifier trained to distinguish LoC from W presented significant accuracy when applied to distinguish N3 from W. The three classifiers presented high and significant (p<0.001) performance when distinguishing W from the brain states used for their training (indicated as self arrows in Fig. 2B): N3 vs.W, <AUC> = 0.948 ± 0.005; LoC vs. W, <AUC> = 0.949 ± 0.004; UWS vs. W, <AUC> = 0.973 ± 0.001 (mean ± std). Next, we used the trained classifiers to sort datasets different from the ones they were originally trained to distinguish. We found that algorithms trained using N3 sleep generalized well to the classification of LoC from W and vice-versa, yielding significant transfer learning (p<0.05) for the classifier trained using N3 sleep and evaluated on LoC (<AUC> = 0.92 ± 0.02), and for the classifier trained using LoC and evaluated on N3 sleep (<AUC> = 0.91 ± 0.01). However, significant transfer learning was not obtained for classifiers trained or evaluated using the UWS dataset, thus establishing that the descriptive classifier distance metric dissociated physiological and pharmacologically-induced states of unconsciousness from the group of patients with most severe disorders of consciousness. This could reflect the different patterns of FC changes that are associated with transient (N3 sleep, LoC) and persistent (UWS) states of unconsciousness (Fig. 2A).

### Regional FC similarity between states of consciousness

Next, we investigated local similarities between states of unconsciousness by means of the connectivity correlation distance. Fig. 3 shows anatomical renderings of the regions presenting a significant correlation of their profiles of FC changes relative to wakefulness between the states indicated in each inset text. For instance, the top left panel highlights the regions whose local FC profile changed similarly during N1 and N2 sleep, in both cases relative to wakefulness. We first observe that most regions in the frontal lobe changed their FC profile similarly during N1 and N2 sleep and that this also happened for most brain regions in the comparison between N2 and N3 sleep, but for fewer regions in the comparison between N1 and N3 sleep. The changes in FC brought upon by N3 sleep were also very similar to those seen in LoC, but their similarity to S was less marked. FC changes during N2 sleep were less similar to LoC than those observed during N3 sleep. Finally, regional FC changes during DoC states (MCS and UWS) were not significantly correlated with those observed during other states of reduced consciousness, and only presented widespread positive correlations between them. Taken together, these results show that states of deeper unconsciousness are more similar between them than compared to transitional states for which consciousness could be partially preserved, such as N1 sleep and S. An exception was found in the comparison between S and LoC, where most brain regions presented similar FC changes, possibly stemming from a similar neurochemical mechanism activated by propofol at different doses. Correlations were more widespread between the states belonging to each different route towards unconsciousness, i.e. between states corresponding to different sleep stages, propofol doses, and DoC severity. As in the results obtained using the classification distance (Fig. 2), DoC behaved very differently from the other states: no significant correlations (|R|>0.5) in regional FC changes were observed between these states and the others.

**Figure 3.**
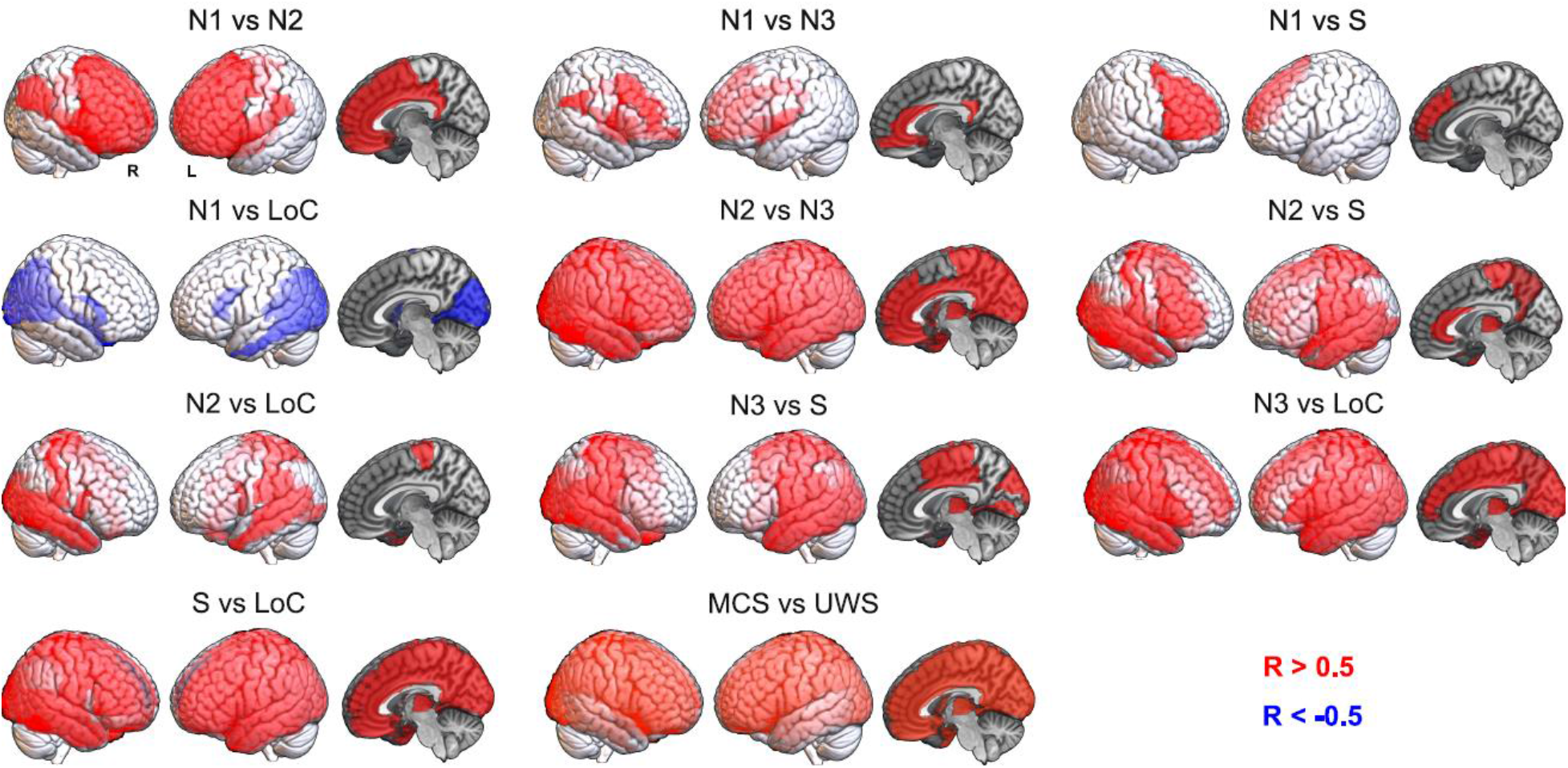
Shared patterns of regional FC changes reflect the progression towards unconsciousness during sleep and propofol-induced anesthesia, but these similarities do not extend to DoC patients. Each panel contains an anatomical rendering of the regions presenting significantly correlated FC changes between the states indicated in the insets. Red indicates a significant positive correlation in regional FC changes (R>0.5), while blue indicates a significant negative correlation (R<-0.5).

### Descriptive distance metrics between states of consciousness

We extended the results shown in Figs. 2 and 3 to include all the possible comparisons between brain states, as well as the comparison based on the similarity of the optimal model parameters (“model parameter distance”, based on the computational model described in Fig. 1). Fig. 4A presents matrices containing z-scores of the aforementioned distance metrics between all pairs of states (note that the classification distance was defined as 1-<AUC>). The matrix elements are presented in graph form in Fig. 4B, with only the top 25% matrix elements being shown.

**Figure 4.**
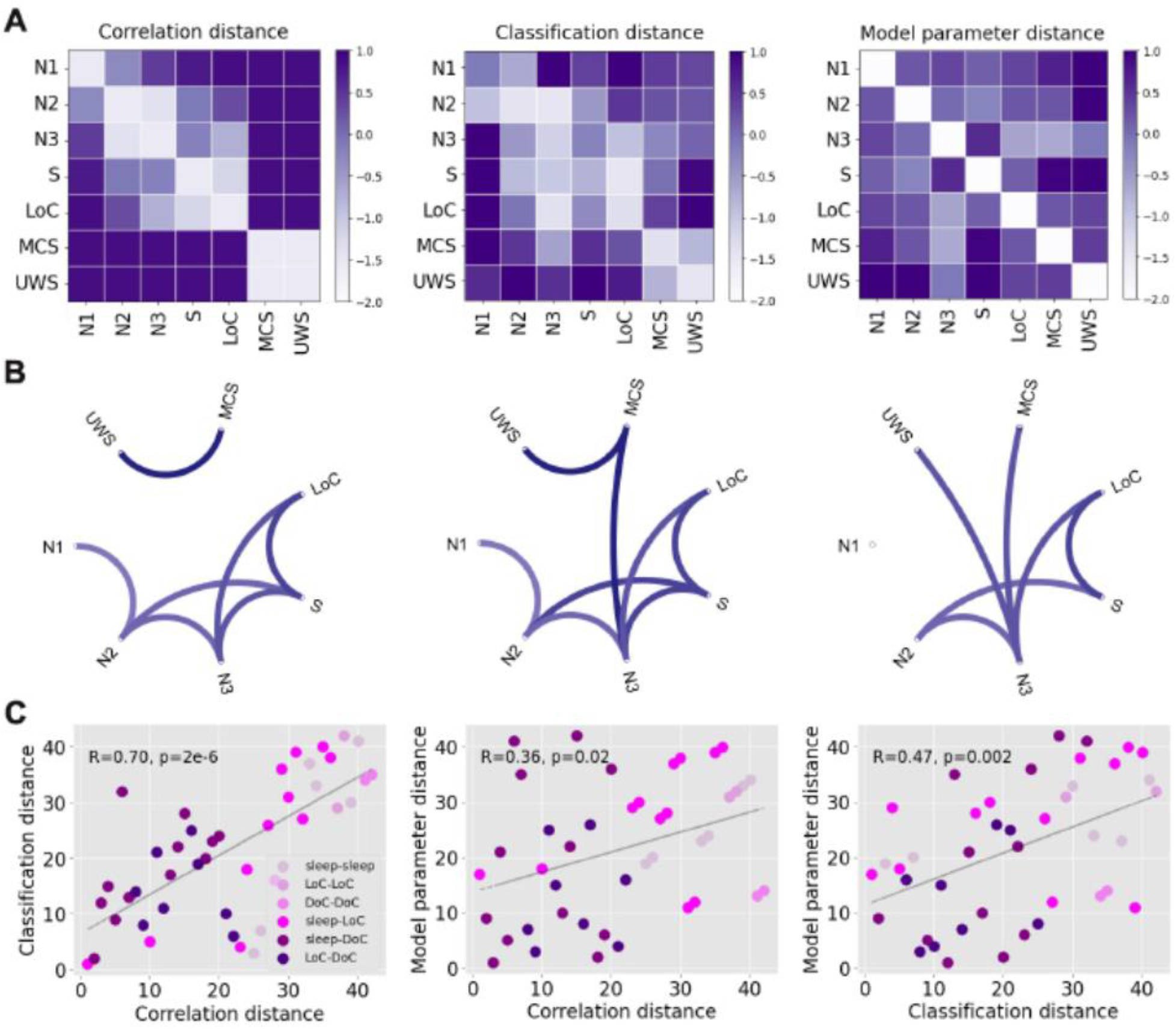
Significant positive correlations between all descriptive distance metrics computed for all pairs of states. (A) Matrices containing z-scores of the correlation, classification and model parameter distances between all pairs of states of consciousness (B) Graph representation of the matrices in panel A, showing only the top 25% matrix elements. (C) Scatterplots establishing the positive and significant correlation between all descriptive distance metrics. Each point represents a pair of states, and the shade of purple indicates one of the following combinations of states: sleep-sleep, LoC-LoC, DoC-DoC, sleep-LoC, sleep-DoC, LoC-DoC. Since variables were converted to ranks prior to the visualization, R and p represent the Spearman’s rank correlation coefficient and its associated p-value, respectively.

The first matrix is based on the proportion of significant regions shown in Fig. 3, i.e., the ratio between the number of significant regions and the 90 regions in the brain atlas (“connectivity correlation distance”). As shown in Fig. 3, contiguous sleep stages presented the lowest distances, while the distance between S/LoC and sleep stages gradually increased from N3 to N1 sleep. As also expected from the previous figures, MCS and UWS patients were highly similar between them but not when compared to the other states. Similar results were obtained for the classification distance (Fig. 4A, second matrix), which is already evident by inspection of the matrix and its associated graph. The results obtained comparing the optimal model parameters (Fig. 4A, third matrix) appear slightly different, but still preserve the three main findings observed for the other metrics: sleep stages of similar depth tended to present the highest similarities, S and LoC were more similar between them than to sleep stages, and DoC patients presented idiosyncratic changes that set them apart from the other states of consciousness.

Fig. 4C shows a quantitative evaluation of the similarity between the three matrices, establishing that each descriptive distance metric can be used to predict all others. In these figures, each point corresponds to a pair of brain states, with X and Y coordinates based on the different combinations of distance metrics. As shown in the last panel, for example, pairs of brain states presenting high machine learning transfer learning accuracy also yielded similar model parameters, and vice-versa. In particular, all three metrics converge in the dissociation between sleep and propofol-induced unconsciousness from DoC patients.

### Perturbational distance between states of consciousness

After optimizing the computational model to reproduce the empirical FC of initial and target states, we systematically simulated the effects of an external periodic perturbation introduced at all pairs of homotopic regions. The perturbation was applied in the model optimized to reproduce the FC of the initial state, and we evaluated how increasing perturbation amplitudes impacted on the model goodness of fit computed relative to the target state. The perturbational distance was computed as the best goodness of fit relative to the target state across amplitudes. According to this definition, a low distance indicates that a suitable combination of perturbation amplitude and stimulation site is capable of displacing the initial FC towards that of the target state, i.e. a transition between initial and target state can be induced by external stimulation in the model.

The perturbational distances between all pairs of physiological, pharmacologically-induced and pathological states of consciousness are summarized in matrix representation in Fig. 5A. A directed graph constructed from these matrix elements is presented in Fig. 5B, where each arrow indicates that certain stimulation parameters induce a transition from the initial to the target state with Δ*GoF* ≤0.3. We observe that certain states receive several directed connections (e.g. W) while others present the opposite behaviour (e.g. N1 sleep). States receiving several connections can be considered stable, since external stimulation easily transitions other states of consciousness towards them, but they generally remain the same when stimulated; conversely, states sending out several connections easily transition into other states when stimulated, and hence can be considered unstable.

**Figure 5.**
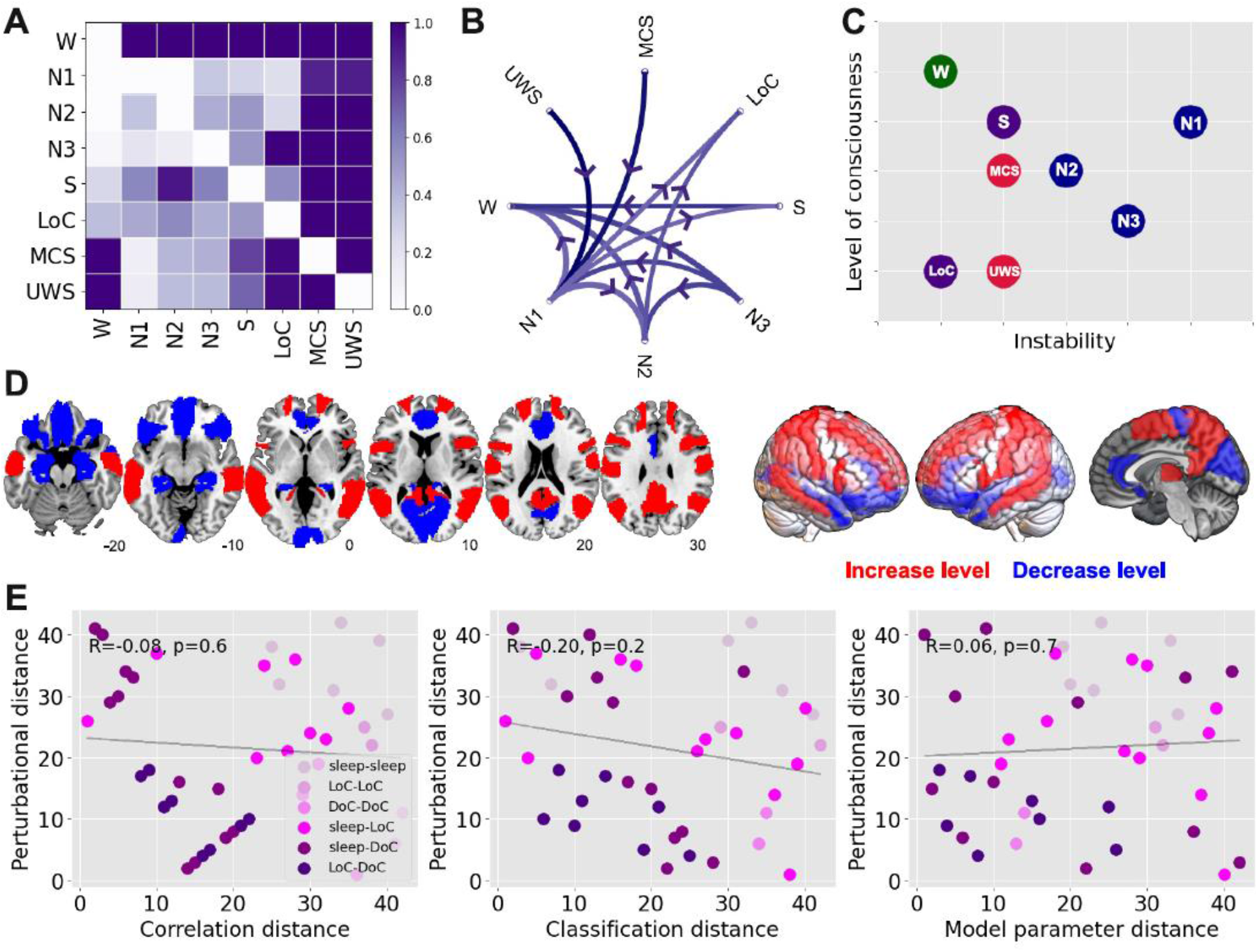
A perturbational metric for the distance between states of consciousness. (A) Matrix representation of the the perturbational distance between all pairs of brain states. (B) Directed graph representation of the matrix in Panel A, showing the possible externally-induced transitions between pairs of states of consciousness (thresholded at Δ*GoF* ≤0.3). (C) Two-dimensional diagram of all states of consciousness according to their level of consciousness (i.e. similarity to wakefulness) and their instability against external perturbations (sleep states are indicated in blue, S and LoC in purple, and DoC in red). (D) Homotopic regions associated with the best Δ*GoF* changes in transitions towards states of reduced consciousness (blue) and increased consciousness (red). (E) Scatter plots illustrating the non-significant correlation between perturbational and descriptive distance metrics. Each point represents a pair of states, and the shade of purple indicates one of the following combinations of states: sleep-sleep, LoC-LoC, DoC-DoC, sleep-LoC, sleep-DoC, LoC-DoC. Since variables were converted to ranks prior to the visualization, R and p represent the Spearman’s rank correlation coefficient and its associated p-value, respectively.

In Fig. 5C we place all states of consciousness into a bi-dimensional diagram according to their similarity to wakefulness (“level of consciousness”, Y-axis) and their instability, defined as the number of outbound connections in Fig. 5B (“instability against perturbations”, X-axis). In this diagram, DoC and LoC appear as stable states of reduced consciousness, while W is both conscious and stable. All sleep stages are comparatively less stable, with N1 sleep being the most fragile against perturbations, consistent with its role as a transitional stage between early and deep sleep. Finally, propofol sedation (S) was intermediate both in conscious level and stability.

We classified transitions in two groups, depending on the initial and target state. One group corresponded to perturbations that increased the level of consciousness (i.e. all states to W, N2/N3 to N1, N3 to N2, LoC to S, MCS to UWS), and another corresponded to perturbations that decreased the level of consciousness (i.e. all reverse transitions). For each pair of states we ranked homotopic regions in terms of their associated optimal Δ*GoF*, and computed the average regional ranking separately across all transitions in the “increase level” and “decrease level” groups. Thus, a high value for a region in the “increase level” group indicates that perturbations applied to that region consistently tended to increase the level of consciousness, and vice-versa for the “decrease level” group. Fig. 5D presents a rendering of the top 50% homotopic regions in each group. Simulated perturbations applied at the bilateral hippocampus, inferior frontal cortex, anterior cingulate cortex and primary visual cortex (calcarine sulcus) systematically resulted in the best Δ*GoF* changes towards states of reduced consciousness. Conversely, perturbations applied at the temporo-parietal junction (bilateral angular gyrus), precuneus, precentral gyrus and middle frontal cortex resulted in the best Δ*GoF* changes towards states of increased consciousness.

In contrast to the results shown in Fig. 4C, the perturbational distance metric provided information complementary to that obtained from the descriptive distance metrics. Fig. 5E shows that the descriptive metrics were not significantly correlated with the perturbational metric; in other words, even though some pairs of states presented similar patterns of FC changes relative to wakefulness, externally-induced transitions between them were forbidden in our computational model.

## Discussion

There are two different but related problems in the study of states of reduced consciousness. The first concerns the identification of such states from a limited amount of behavioral information and non-invasive brain activity recordings. This is the challenge faced by clinicians in the identification of DoC, a difficult task with up to 40% consensus-based misdiagnosis rate (32), as well as by anesthesiologists in the detection and prevention of intraoperative awareness (33). The second problem concerns the manipulation of conscious states by means of externally induced perturbations, either to induce unconsciousness (i.e. anesthesia) or to restore conscious wakefulness in patients (34–3) These two problems map onto the dimensions we explored in the present work and are summarized in Fig. 5C. Previous data-driven and theoretical developments for the detection of consciousness from neuroimaging data (10–13,15) represent partial solutions: what is also needed is an exhaustive and systematic method to investigate the potential behaviour of global brain states under external stimulations. We pursued this approach by combining different sources of empirical information with simple but conceptually rich models of whole-brain activity, which allowed us to explore the stability of different states of consciousness from passive recordings of fMRI data. Importantly, we showed that the perturbational analysis provided information complementary to the results of statistical and machine learning techniques applied directly to the data.

The representation of whole-brain activity by coupled dynamical systems was a crucial step in our analysis. While previous experimental studies investigated the effects of localized external perturbations during states of reduced awareness in humans (34–36,39,40), the systematic exploration of targeted stimulation is possible by the freedom granted by computational models. We opted to allow regional variability in the bifurcation parameters of the model, since different brain regions could be more or less relevant to induce transitions between certain states of consciousness, and this variability could depend upon the proximity of regional dynamics to the bifurcation point (28,29). The use of RSNs to constrain this regional variability, combined with other sources of empirical data, increased the conceptual interpretability of our computational model. Due to the semi-empirical nature of the model, we were able to show that the inferred parameters reflected the similarities between states of consciousness observed in FC patterns; for instance, the distance metric based on random forest transfer learning accuracy was significantly correlated with the metric obtained from the comparison of the optimal model parameters (Fig. 4C). However, the addition of external stimulation to the model resulted in a distance metric that was independent from those obtained without perturbational analysis (Fig. 5E). In particular, this distance metric could not be predicted by the similarity of the underlying model parameters, suggesting that perturbations are amplified by the system nonlinearities, a behaviour characteristic of systems posed at or near a dynamical instability (26,29,41).

States of consciousness can be analyzed in functional terms (i.e. behavior and cognition) as well as by quantitative metrics derived from brain activity measurements. It is becoming increasingly clear that functional analysis framed in terms of unidimensional “levels of consciousness” could be insufficient to capture the richness and heterogeneity of conscious states (5). A conceptually similar unidimensional characterization is also pursued by different quantitative indices computed from neural activity recordings, such as information integration (42), compressibility (15,25), causal density (18), the perturbational complexity index (PCI) (12), and other data-driven metrics. Some of the most commercially successful markers of conscious awareness, such as the bispectral index for anesthesia monitoring (10), are based on this approach. Our work questions whether markers of this kind can be sufficient, since they do not address the potential behavior of the system against perturbations. Here, it is important to draw a distinction between the stability of ongoing neural dynamics and the stability of global brain states. PCI (12) and other related data-driven techniques (43) estimate the response of within-state activity to perturbations, but it is assumed that the applied perturbation does not result in a transition between brain states. In contrast, our interest lies precisely in determining the likelihood of observing such a transition upon external stimulation. Clearly, these different approaches can yield independent and complementary information, since our perturbational distance metric showed a dissociation between general anesthesia and deep sleep, while both states present comparable PCI values (12). Finally, the results obtained from our perturbational analysis are fully consistent with well-understood differences in responsiveness between sleep and propofol-induced anesthesia; for example, with the observation that arousals are more easily elicited during sleep than under the effects of anesthesia (equivalently, surgeons cannot wait for the onset of sleep to start operating). While responsiveness can be probed by direct sensory stimulation, our model was capable of reproducing the same result through the exhaustive exploration of all pairs of homotopic regions, i.e. not restricted to sensory regions.

Deep sleep and propofol-induced anesthesia present similarities in terms of brain activity and their associated neurochemical pathways. Both states are associated with slow (44) and regular activity of cortical origin (15,25), breakdown of large-scale FC (45,46), and increased inhibitory neurotransmission (47); furthermore, propofol anesthesia may result in sleep-like homeostatic regulation (48). The results we presented in Fig. 2 are in line with these observations and also point towards marked differences between these states and the conditions of MCS and UWS patients, which have been previously shown to present distinct changes in EEG dynamics (49). A recent article demonstrated significant transfer learning between datasets comprising propofol anesthesia and DoC patients, which is at odds with the results of our analysis (50). We believe this contradiction could arise due to the large variability that exists between cohorts of brain-injured patients (51). Future studies should attempt to settle this issue by investigating larger and more homogenous patient populations.

Within the different stages of human sleep, we showed that N1 sleep (a transitional stage between wakefulness and deep sleep) presented the highest instability against perturbations. N1 sleep is characterized by diminished thalamo-cortical coupling with preserved cortical activation, a condition compatible with conscious mentation and imagery during the onset of sleep (52). From an evolutionary perspective, it is reasonable that N1 sleep is susceptible to transitioning towards wakefulness upon external stimulation, since during this stage the individual is vulnerable to environmental threats, while offline information processing associated with learning and memory consolidation is not yet taking place (53). On the other hand, propofol-induced unconsciousness and DoC are either artificially induced or arise as a consequence of brain injury, and thus do not reflect the adaptive pressures that constrain the stability of human sleep. These constraints are also reflected in the sequence of states that comprise human sleep: the orderly progression from wakefulness to N3 sleep (54) is disrupted when sudden awakenings occur. The output of our model was consistent with these dynamics, since external perturbation could only induce transitions from N3 sleep to wakefulness, but failed to elicit similar transitions towards intermediate sleep stages.

The non-reversible nature of severe DoC could be linked to alterations in the underlying structural connectivity of the brain, as a consequence of injury (55,56). While whole-brain functional connectivity is known to be preserved even during states of deep unconsciousness (57,58), it tends to reduce towards the structural connectivity backbone (20,59,60), suggesting that this backbone imposes a limit to the functional disintegration that is possible in healthy brains. However, the analysis of the changes reported in Fig. 2 reveals that patients present a substantially different functional architecture compared to that seen during sleep and propofol-induced loss of consciousness, which could stem from a fundamentally different (and more variable) organization of anatomical connections. We note that our analysis does not preclude the possibility of inducing transitions towards conscious wakefulness in patients, but individually chosen targets might be necessary due to the aforementioned variability. Also, it could be possible that the recovery of consciousness can be accelerated by neurochemical and pharmacological means (61), which cannot be easily accommodated within the proposed modeling framework.

The choice of periodic stimulation was determined mainly by the local dynamics, which consisted of nonlinear oscillators with a single natural frequency. However, future modeling efforts incorporating more complex dynamics could allow *in silico* rehearsal of interventions with ampler neurobiological interpretation; for instance, a dynamic mean-field model informed by empirical receptor density maps could be used to explore the result of activating specific neurotransmitter systems (e.g. serotonin, dopamine) (62,63).This flexibility could be used to extend our analysis to other conscious states, such as those seen in certain psychiatric patients. That different psychiatric conditions can present distinct levels of stability is known to clinical practitioners who have encountered patients suffering from bipolar disorder on one extreme, and catatonic patients on the other. Also, the application of machine learning classifiers combined with computational models could inform the hypothesis that certain pharmacological interventions mimic the symptomatology of certain psychiatric syndromes, such as in the psychotomimetic hypothesis of serotonin 2A receptor agonists (also known as “psychedelics”) (64).

Our results should not only be discussed in terms of the allowed transitions between states, but also in terms of which regions are associated with those transitions, and how those transitions depend on the external forcing amplitude. In Fig. 5D we showed that transitions towards states of heightened consciousness were systematically linked to perturbations located in the precuneus, temporo-parietal junction and the middle frontal cortex, regions presenting a significant overlap with the default mode network (65) and in line with electrical stimulation targets shown to improve behavioural signatures of consciousness in DoC patients (38). These regions have also been shown to robustly reflect the level of consciousness (46,55,66,67) and conscious information access in cognitive neuroscience paradigms (1). Consistently with previous work (28), the effect of perturbing these regions also depended on the amplitude of the periodic forcing term; for instance, perturbations applied to the temporo-parietal junction asymptotically increased the similarity to wakefulness, while a small perturbation located at the prefrontal cortex sufficed to reproduce an arousal. The qualitatively different behavior upon simulated perturbations represents a set of rich predictions to be addressed by future experiments.

While each independent source of empirical information incorporated into our model increased its interpretability, it also imposed specific limitations to our analysis. For instance, functional connectivity was estimated from recordings acquired in different centers, which could represent a potential source of confounds. However, since our approach focused on transfer learning accuracy and each machine learning classifier was trained using a control group acquired using a matched scanner and data acquisition protocol, we believe it was less vulnerable to classification biases related to different experimental conditions. Also, the use of anatomical connectivity estimated in a group of healthy participants could represent a limitation for the modeling of patient data. Nevertheless, since brain-injured patients may present heterogeneous lesion locations (51) it could be that the average healthy connectivity constitutes a reasonable first estimate. Also, since the perturbations failed to induce widespread transitions between LoC/sleep and DoC (even when considering the same anatomical connectivity) we can expect that this result will be furthered when incorporating more accurate group-specific connectivity.

In conclusion, the investigation of dynamical stability can be informative for the characterization of different brain states, allowing the dissociation between reversible vs. non-reversible and pharmacological vs. physiological states, with potential applications to neurologic and psychiatric conditions associated with persistent states of abnormal consciousness and cognition. We expect that future metrics to monitor levels of sleep, anesthesia and residual consciousness in brain injured patients are expanded to represent this additional dimension, with positive consequences in clinical practice and in the neuroscientific investigation of human consciousness and its disorders.

## Materials and Methods

### Experimental data

We analyzed fMRI recordings from 81 participants scanned at two independent research sites: Frankfurt: 15 subjects during wakefulness and sleep; Liège: 14 healthy subjects during wakefulness and under propofol sedation and anesthesia; 16 patients diagnosed as MCS, 15 patients diagnosed as UWS, and 21 healthy and awake controls.

#### Sleep dataset

Simultaneous fMRI and EEG was measured for a total of 73 subjects (written informed consent, approval by the local ethics committee) in Frankfurt (Germany). EEG via a cap (modified BrainCapMR, Easycap, Herrsching, Germany) was recorded continuously during fMRI acquisition (1505 volumes of T2*-weighted echo planar images, TR/TE = 2080 ms/30 ms, matrix 64 × 64, voxel size 3 × 3 × 2 mm^3^, distance factor 50%; FOV 192 mm2) with a 3 T Siemens Trio (Erlangen, Germany). An optimized polysomnographic setting was employed (chin and tibial EMG, ECG, EOG recorded bipolarly [sampling rate 5 kHz, low pass filter 1 kHz] with 30 EEG channels recorded with FCz as the reference [sampling rate 5 kHz, low pass filter 250 Hz]. Scalp potentials measured with EEG allow the classification of sleep into 4 stages (wakefulness, N1, N2 and N3 sleep) according to the American Academy of Sleep Medicine (AASM) rules (54). Pulse oximetry and respiration were recorded via sensors from the Trio [sampling rate 50 Hz]) and MR scanner compatible devices (BrainAmp MR+, BrainAmpExG; Brain Products, Gilching, Germany), facilitating sleep scoring during fMRI acquisition. We selected 15 subjects who reached stage N3 sleep (deep sleep) and contiguous time series of least 200 volumes for all sleep stages. Previous publications based on this dataset can be consulted for further details (see,e.g., ref. 68).

#### Propofol sedation and anesthesia

Resting-state fMRI volumes from 18 healthy subjects were acquired in four different states following propofol injection: wakefulness, sedation, unconsciousness, and recovery. Data acquisition was performed in Liège (Belgium) (written informed consent, approval by the Ethics Committee of the Medical School of the University of Liège). Subjects fasted for at least 6 h from solids and 2 h from liquids before sedation. During the study and the recovery period, electrocardiogram, blood pressure, pulse oximetry (SpO2), and breathing frequency were continuously monitored (Magnitude 3150M; Invivo Research, Inc., Orlando,FL). Propofol was infused through an intravenous catheter placed into a vein of the right hand or forearm. An arterial catheter was placed into the left radial artery. Throughout the study, the subjects breathed spontaneously, and additional oxygen (5 l/min) was given through a loosely fitting plastic facemask. The level of consciousness was evaluated clinically throughout the study with the scale used in ref. 69. The subject was asked to strongly squeeze the hand of the investigator. She/he was considered fully awake or to have recovered consciousness if the response to verbal command (“squeeze my hand”) was clear and strong (Ramsay 2), as sedated if the response to verbal command was clear but slow (Ramsay 3), and as unconscious, if there was no response to verbal command (Ramsay 5–6). Ramsay scale verbal commands were repeated twice for each consciousness level assessment. Functional MRI acquisition consisted of resting-state functional MRI volumes repeated in the four states: normal wakefulness (Ramsay 2), sedation (Ramsay 3), unconsciousness (Ramsay 5), and recovery of consciousness (Ramsay 2). The typical scan duration was half an hour for each condition, and the number of scans per session (200 functional volumes) was matched across subjects to obtain a similar number of scans in all states. Functional images were acquired on a 3 Tesla Siemens Allegra scanner (Siemens AG, Munich, Germany; Echo Planar Imaging sequence using 32 slices; repetition time = 2460 ms, echo time = 40 ms, field of view = 220 mm, voxel size = 3.45×3.45×3 mm^3^, and matrix size = 64×64×32). Previous publications based on this dataset can be consulted for further details (see, e.g., ref. 46).

#### Disorders of consciousness

The dataset comprised resting-state fMRI volumes on healthy controls (>18 years old and free of psychiatric and neurological history) and unsedated patients presenting disorders of consciousness (Department of Radiology, Centre Hospitalier Universitaire (CHU), Liège) (written informed consent to participate in the study was obtained directly from healthy control participants and the legal surrogates of the patients, approval by the Ethics Committee of the Medical School of the University of Liège). The cohort included 21 healthy controls (8 females; mean age, 45 ± 17 years), 43 patients (25 in MCS, 18 in UWS, 12 females; mean age, 47 ± 18 years. See Supplementary Information with single subject demographic information). UWS patients show signs of preserved vigilance, but do not exhibit non-reflex voluntary movements, and are incapable of establishing functional communication (70). Patients in MCS show more complex behavior indicative of awareness, such as visual pursuit, orientation response to pain, and non-systematic command following; nevertheless, these signs are consistent but may be manifested sporadically (71). The inclusion criteria for patients were brain damage at least 7 days after the acute brain insult and behavioral diagnosis of MCS or UWS performed with the Coma Recovery Scale–Revised (CRS-R) (9). The CRS-R is currently the most sensitive scale to characterize disorders of consciousness and evaluates and includes 23 arranged items organized on subscales for auditory, visual, motor, oromotor, communication, and arousal function. Each item assesses the presence or absence of specific physical signs, which represent the integrity of brain function as presence or absence of cognitively mediated responsiveness.

Data were acquired on a 3T Siemens TIM Trio MRI scanner (Siemens Medical Solutions, Erlangen, Germany): 300 T2*-weighted images were acquired with a gradient-echo echo-planar imaging (EPI) sequence using axial slice orientation and covering the whole brain (32 slices; slice thickness, 3 mm; repetition time, 2000 ms; echo time, 30 ms; voxel size, 3 × 3 × 3 mm; flip angle, 78°; field of view, 192 mm by 192 mm). A structural T1 magnetization-prepared rapid gradient echo (MPRAGE) sequence (120 slices; repetition time, 2300 ms; echo time, 2.47 ms; voxel size, 1.0 × 1.0 × 1.2 mm; flip angle, 9°) (Demertzi et al., 2019)

#### fMRI preprocessing

For each participant and for each brain state, we used FSL tools to extract and average the BOLD signals from all voxels. The FSL preprocessing included a 5mm spatial smoothing (FWHM), bandpass filtering between 0.01-0.1 Hz, and brain extraction (BET), followed by a transformation to a standard space (2mm MNI brain) and down sampling for a final representation in a 45×54×45, 2mm voxel space.

The following preprocessing steps were performed using specially developed Matlab scripts. First, we correct the data by performing regressions between the displacement parameters and the average signal and its first derivative, and eliminate these correlations, since they mainly inform movement. Continuing, we perform a *scrubbing* process by which we eliminate subjects that present significant movements in more than 20% of the temporal frames recorded, with a criterion for movement significance set as an absolute displacement between consecutive frames larger than 0.5 mm. For the remaining subjects, we removed the first 3 frames and those which exceed the aforementioned threshold. Finally, we averaged all voxels within each ROI defined in the automated anatomical labeling (AAL) atlas, considering only the 90 cortical and subcortical non-cerebellar brain regions (72) to obtain one BOLD signal per ROIs.

This way, we obtain datasets with comparable smoothness and stability that can be compared across conditions.

During preprocessing, 4 subjects were removed from the anesthesia data set, as well as 9 MCS patients and 3 UWS patients. (See Supplementary Information)

#### Structural Connectivity

The structural connectome was obtained applying diffusion tensor imaging (DTI) to diffusion weighted imaging (DWI) recordings from 16 healthy right-handed participants (11 men and 5 women, mean age: 24.75 ± 2.54 years) recruited online at Aarhus University, Denmark. For each participant a 90×90 SC matrix was obtained that represents the connectivity between ROIs. Data preprocessing was performed using FSL diffusion toolbox (Fdt) with default parameters. The probtrackx tool in Fdt was used to provide automatic estimation of crossing fibers within each voxel, which has been shown to significantly improve the tracking sensitivity of non-dominant fiber populations in the human brain. The connectivity probability from a seed voxel *i* to another voxel *j* was defined as the proportion of fibers passing through voxel *i* that reached voxel *j* (sampling of 5000 streamlines per voxel [73]). All the voxels in each AAL parcel were seeded (i.e. grey and white matter voxels were considered). The connectivity probability *P_ij_* from region *i* to region *j* was calculated as the number of sampled fibers in region *i* that connected the two regions, divided by 5000 × n, where n represents the number of voxels in region *i*. The resulting SC matrices were computed as the average across voxels within each ROI in the AAL thresholded at 0.1% (i.e. a minimum of five streamlines) and normalized by the number of voxels in each ROI. Finally, the data were averaged across participants.

### Multivariate machine learning classifiers

We trained random forest classifiers (74) to distinguish reduced states of consciousness from wakefulness based on empirical individual FC matrices (fully connected weighted matrices computed using Pearson’s linear correlation coefficient between BOLD time series from each subject), using a five-fold crossvalidation procedure to estimate classifier accuracy. Classifiers were first trained to distinguish between wakefulness and a state of reduced consciousness, and their accuracy was then tested in the classification between wakefulness and all other states of consciousness (i.e. transfer learning accuracy was assessed). Random forest classifiers were trained using scikit-learn (https://scikit-learn.org/) (75). We trained random forest classifiers with 1000 decision trees and a random subset of features of size equal to the (rounded) square root of the total number of features. The quality of each split in the decision trees was measured using Gini impurity, and the individual trees were expanded until all leaves were pure (i.e. no maximum depth was introduced). No minimum impurity decrease was enforced at each split, and no minimum number of samples was required at the leaf nodes of the decision trees (the classifier hyperparameters can be found in https://scikit-learn.org/).

To assess the statistical significance of the classifier accuracy values, we trained and evaluated a total of 1000 random forest classifiers using the same features (i.e. FC matrices) as inputs, but scrambling the class labels. We then constructed an empirical p-value by counting how many times the accuracy of the classifier with scrambled class labels was greater than that of the original classifier All accuracies were computed as the area under the receiver operating characteristic curve (AUC) and considered significant at p<0.05. Subsequently, the generalizability of the classifiers to distinguish other sleep states from wakefulness was evaluated by applying both the original and scrambled classifiers, and constructing p-values analogously.

### Regional FC similarity

For all states of consciousness, we computed the average functional connectivity (FC) of each AAL region and subtracted the average FC computed from the wakefulness data, thus yielding a regional profile of FC changes for each state. We then computed the connectivity correlation distance between pairs of states S_1_, S_2_ with FC matrices C_1_, C_2_ as follows,

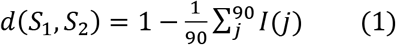

Where I(j) is defined as,

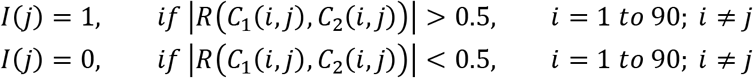

Here, the FC of state 1 is obtained as the average across subjects with wakefulness subtracted, and analogously for state 2.

### Whole-brain model

We implemented a whole-brain model consisting of a network of nonlinear oscillators coupled by the structural connectome (SC). Each oscillator was modeled by a normal form of a Hopf bifurcation and represented the dynamics of one of the 90 brain regions in the AAL template. The key neurobiological assumption is that dynamics of macroscopic neural masses can range from fully synchronous (i.e. activated state with self-sustained oscillations) to a stable asynchronous state governed by random fluctuations, with an intermediate state presenting complex temporal features linked to noise-induced transitions through the bifurcation point (28,29). A secondary assumption is that fMRI can capture the dynamics from both regimes with sufficient fidelity to be modeled by the equations.

Without coupling, the local dynamics of brain region j are modeled by the complex-valued equation,

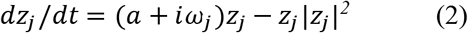

In this equation *z_j_* is a complex-valued variable (*z_j_* = *x_j_* + *iy_j_*), and *ω_j_* is the intrinsic oscillation frequency of node *j*. The intrinsic frequencies ranged from 0.04-0.07 Hz and were determined by the averaged peak frequency of the bandpass-filtered fMRI signals of each individual brain region. The parameter **a** is known as the bifurcation parameter and controls the dynamical behavior of the system. For **a**<0 the phase space presents a unique stable fixed point at *z_j_* = 0, thus the system asymptotically decays towards this point. For **a**>0 the stable fixed point changes its stability, giving rise to a limit cycle and to self-sustained oscillations with frequency *f_j_* = *ω_j_*/*2π* and amplitude proportional to the square root of **a** (see Fig. 1).

The coordinated dynamics of the resting state activity are modeled by introducing coupling determined by the SC. Nodes *i* and *j* are coupled by *C_ij_* (the *i,j* entry of the SC matrix). To ensure oscillatory dynamics for **a**>0, the SC matrix was scaled to a maximum of 0.2 (weak coupling condition) (29). In full form, the coupled differential equations of the model are the following,

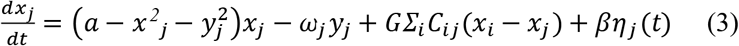

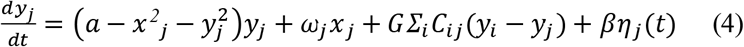

The parameter G represents a global coupling factor that scales SC equally for all the nodes. These equations were integrated to simulate empirical fMRI signals using the Euler-Maruyama algorithm with a time step of 0.1 seconds. *η*_j_ represents additive Gaussian noise in each node and scaled by factor β fixed at 0.04. When **a** is close to the bifurcation (**a**~0) the additive Gaussian noise gives rise to complex dynamics as the system continuously switches between both sides of the bifurcation.

### Fitting to empirical data

We selected the group-averaged static functional connectivity of each state of consciousness as the empirical observable to be fitted by the model. The BOLD signals corresponding to each ROI in the AAL template were filtered in the frequency range of 0.04–0.07 Hz, since this frequency band has been shown to contain more reliable and functionally relevant information compared to other frequency bands, and also to be less affected by noise (76–79). Subsequently, the filtered time series were transformed into z-scores. For each state of consciousness, the amount of participants was selected based on the presence of uninterrupted epochs of that state lasting more than 194 samples (W_sleep_= 15; N1=15; N2=15; N3=15; W_prop_=14; S=14; LoC=14;W_con_=21; MCS:16; UWS:15). Afterwards, the FC matrix was computed as the matrix of Pearson’s correlation coefficients between the BOLD signals of all pairs of regions of interest (ROIs) in the AAL template. Fixed-effect analysis was used to obtain group-level FC matrices, meaning that Fisher’s R-to-z transform z=atanh(R) was applied to the correlation values before averaging over participants within each state of consciousness.

We applied the model described by Eqs. 3 and 4 to simulate BOLD signals for each ROI. We used an anatomical prior based on six RSNs to constrain how different groups of nodes could contribute independently to the final bifurcation parameters. Each local bifurcation parameter was obtained as the linear combination of the contribution of the RSNs spanning that ROI. In this way, we embedded the parameters governing the dynamics of the 90 ROIs into a six-dimensional parameter space defined by the independent contributions of the RSNs (28). We simulated the same number of samples for each subject and the same number of subjects per state, and then we followed a procedure to compute the simulated FC identical to the one used for the empirical data. We used the structure similarity index (SSIM) (Wang et al., 2004) as a metric to compare the simulated and empirical FC, thus defining the goodness of fit (GoF) for parameter optimization (the target fitting function was defined as 1-GoF). We implemented a genetic algorithm to optimize the six parameters and maximize the GoF of the model. For each state of consciousness, we simulated an initial population of 10 elements, 200 generations of offspring and then we performed 100 independent runs of the genetic algorithm. Previous work implementing the same optimization procedure can be consulted for further details (28). Finally, we selected the combination of parameters yielding the simulated FC with the lowest GoF among the 100 runs of the algorithm.

### Perturbational distance

The external perturbation was represented as an additive periodic forcing term incorporated to the equation of each node, given by *F_j_* = *F*_0_*j*__*cos*(*ω_j_t*), where *F*_0_*j*__ is the perturbation amplitude and *ω_j_* is the natural frequency of node *j*, computed directly from the BOLD time series. The effects of the perturbation were investigated systematically for all 45 pairs of homotopic regions in the AAL atlas, with the purpose of providing a conceptual model of the effects of transcranial alternating current stimulation (tACS). This perturbation was initially applied in the model with parameters chosen to reproduce an initial state, and the amplitude (*F*_0_*j*__) of node *j* and its homotopic pair was parametrically increased from 0 to 2 in steps of 0.1 (averaging 100 independent simulations for each node pair and *F*_0_*j*__ value). For each value of *F*_0_*j*__ the resulting FC matrix was computed, and its similarity to the FC of the target state was determined as follows,

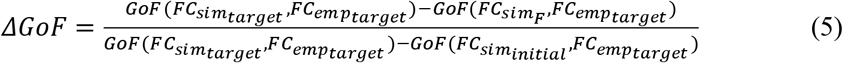

In this equation, *FC_sim_F__* is the FC matrix obtained with the perturbation, *FC_emp_target__* represents the empirical FC matrix of the target state, *FC_sim_target__* is the simulated FC matrix of the target state, and *FC_sim_initial__* is the simulated matrix of the initial state. According to this normalization, as Δ*GoF* approaches 0 the simulation with optimal bifurcation parameters for the initial state plus the perturbation approaches the best empirical fit of the model to the target FC. Conversely, as Δ*GoF* increases the perturbation fails to change the FC in the direction of the optimal FC of the target state (28).

## Acknowledgments

The study was supported by the University and University Hospital of Liège, the Belgian National Funds for Scientific Research (FRS-FNRS), the European Union’s Horizon 2020 Framework Programme for Research and Innovation under the Specific Grant Agreement No. 945539 (Human Brain Project SGA3), the European Space Agency (ESA) and the Belgian Federal Science Policy Office (BELSPO) in the framework of the PRODEX Programme, “Fondazione Europea di Ricerca Biomedica”, the Bial Foundation, the Mind Science Foundation and the European Commission, the fund Generet, the King Baudouin Foundation, DOCMA project [EU-H2020-MSCA–RISE–778234]. SL is research director at FRS-FNRS. AI is partially supported by grants from CONICET, FONCyT-PICT 2017-1818, FONCyT-PICT 2017 1820, ANID/FONDAP/15150012, the Interamerican Development Bank (IDB), GBHI ALZ UK-20-639295, NIH NIA R01 AG057234, and the MULTI-PARTNER CONSORTIUM TO EXPAND DEMENTIA RESEARCH IN LATIN AMERICA (ReDLat). The contents of this publication are solely the responsibility of the authors and do not represent the official views of these Institutions. ET is supported by funding from Agencia Nacional De Promocion Cientifica Y Tecnologica (Argentina), grant PICT-2018-03103. The authors acknowledge the Toyoko 2020 program for granting cloud computing services.The authors thank the patients and their families for participation in the study, as well as the whole staff from the Radiodiagnostic department at University Hospital of Liège. We are highly grateful to the members of the Liège Coma Science Group for their assistance in clinical evaluations.

## Supplementary Information

**Table S1.** Patients’ demographic and clinical characteristics. **Diagnosis**: MCS: minimally conscious state, UWS: vegetative state/ unresponsive wakefulness syndrome. **Etiology**: 1: traumatic brain injury, 2: anoxia, 3: other **Coma Recovery Scale-Revised subscales:Auditory function** 4: Consistent Movement to Command, 3: Reproducible Movement to Command, 2: Localization to Sound, 1: Auditory Startle, 0: None. **Visual function** 5: Object Recognition, 4: Object Localization: Reaching, 3: Visual Pursuit, 2: Fixation, 1: Visual Startle, 0: None. **Motor function** 6: Functional Object Use, 5: Automatic Motor Response, 4: Object Manipulation, 3: Localization to Noxious Stimulation, 2: Flexion Withdrawal, 1: Abnormal Posturing, 0: None/Flaccid. **Oromotor/Verbal function** 3: Intelligible Verbalization, 2: Vocalization/Oral Movement, 1: Oral Reflexive Movement, 0: None. **Communication scale** 2: Functional: Accurate, 1: Non-Functional: Intentional, 0: None. **Arousal scale** 3: Attention, 2: Eye Opening without stimulation, 1: Eye Opening with stimulation 0: Unarousable. Inclusion field stands for the subject that were included in the full analysis after fMRI pre-processing.

| Diagnosis | Gender | Age at Scan | Etiology | Days since Injury | Comma Recovery Scale-Revised |  |  |  |  |  | #CRS-R assessment | Included |
| --- | --- | --- | --- | --- | --- | --- | --- | --- | --- | --- | --- | --- |
|  |  |  |  |  | Aud. Function | Vis. Function | Mot. Function | Oro./V<br>er<br>Function | Communication | Aurosa<br>l |  |  |
| MCS | F | 34 | 1 | 3034 | 3 | 3 | 2 | 2 | 0 | 2 | 5 | yes |
| MCS | M | 62 | 2 | 13 | 0 | 3 | 2 | 1 | 0 | 1 | 6 | yes |
| MCS | F | 59 | 3 | 21 | 1 | 3 | 2 | 0 | 0 | 2 | 7 | yes |
| MCS | M | 30 | 1 | 246 | 3 | 2 | 2 | 2 | 0 | 1 | 3 | yes |
| MCS | M | 83 | 3 | 13 | 3 | 0 | 2 | 1 | 0 | 0 | 5 | yes |
| MCS | M | 34 | 3 | 1077 | 3 | 5 | 2 | 0 | 1 | 1 | 4 | yes |
| MCS | M | 52 | 3 | 20 | 3 | 3 | 2 | 2 | 1 | 2 | 4 | no |
| MCS | M | 47 | 1 | 533 | 3 | 5 | 2 | 1 | 0 | 2 | 13 | yes |
| MCS | M | 38 | 3 | 1854 | 1 | 3 | 2 | 2 | 0 | 1 | 4 | yes |
| MCS | M | 29 | 3 | 64 | 1 | 3 | 2 | 1 | 0 | 2 | 2 | yes |
| MCS | M | 41 | 2 | 9900 | 3 | 3 | 2 | 2 | 0 | 2 | 8 | no |
| MCS | M | 23 | 1 | 301 | 1 | 3 | 2 | 1 | 0 | 2 | 6 | yes |
| MCS | M | 53 | 2 | 1241 | 0 | 3 | 2 | 2 | 0 | 2 | 5 | no |
| MCS | F | 46 | 3 | 242 | 1 | 3 | 2 | 1 | 0 | 1 | 7 | yes |
| MCS | M | 60 | 1 | 15 | 3 | 5 | 5 | 3 | 1 | 1 | 1 | no |
| MCS | M | 25 | 1 | 1157 | 3 | 5 | 5 | 1 | 0 | 2 | 5 | no |
| MCS | M | 61 | 1 | 135 | 3 | 4 | 2 | 1 | 0 | 2 | 7 | yes |
| MCS | M | 68 | 1 | 360 | 0 | 3 | 2 | 0 | 0 | 2 | 3 | yes |
| MCS | M | 35 | 1 | 1331 | 3 | 0 | 2 | 1 | 0 | 2 | 10 | no |
| MCS | F | 48 | 1 | 291 | 2 | 1 | 2 | 2 | 0 | 1 | 8 | no |
| MCS | M | 73 | 3 | 35 | 0 | 2 | 0 | 1 | 0 | 1 | 3 | yes |
| MCS | M | 23 | 1 | 645 | 3 | 3 | 0 | 1 | 0 | 2 | 8 | yes |
| MCS | M | 11 | 2 | 1482 | 3 | 4 | 2 | 2 | 1 | 2 | 9 | yes |
| MCS | F | 48 | 2 | 215 | 3 | 0 | 5 | 2 | 3 | 0 | 9 | no |
| MCS | M | 66 | 1 | 674 | 3 | 0 | 1 | 2 | 3 | 0 | 5 | no |
| UWS | F | 52 | 3 | 283 | 1 | 0 | 2 | 2 | 0 | 1 | 8 | yes |
| UWS | M | 36 | 2 | 6709 | 1 | 0 | 2 | 1 | 0 | 2 | 6 | yes |
| UWS | M | 30 | 2 | 743 | 1 | 0 | 2 | 1 | 0 | 2 | 7 | yes |
| UWS | M | 74 | 2 | 92 | 1 | 0 | 1 | 1 | 0 | 1 | 9 | yes |
| UWS | F | 44 | 2 | 8 | 0 | 0 | 1 | 0 | 0 | 1 | 2 | yes |
| UWS | M | 67 | 3 | 43 | 1 | 0 | 2 | 1 | 0 | 1 | 5 | yes |
| UWS | M | 63 | 2 | 30 | 1 | 1 | 2 | 2 | 0 | 2 | 1 | yes |
| UWS | M | 31 | 1 | 849 | 1 | 1 | 2 | 1 | 0 | 2 | 11 | yes |
| UWS | M | 65 | 2 | 26 | 0 | 0 | 1 | 0 | 0 | 1 | 2 | yes |
| UWS | M | 87 | 3 | 7 | 0 | 0 | 2 | 1 | 0 | 1 | 1 | no |
| UWS | F | 41 | 2 | 1572 | 1 | 0 | 1 | 1 | 0 | 2 | 8 | yes |
| UWS | F | 50 | 2 | 38 | 0 | 0 | 0 | 2 | 0 | 1 | 3 | yes |
| UWS | M | 44 | 2 | 27 | 1 | 0 | 1 | 0 | 0 | 2 | 1 | yes |
| UWS | F | 16 | 2 | 27 | 1 | 0 | 1 | 1 | 0 | 1 | 6 | no |
| UWS | F | 49 | 2 | 129 | 1 | 0 | 0 | 1 | 0 | 2 | 5 | yes |
| UWS | M | 36 | 2 | 2031 | 1 | 0 | 1 | 2 | 0 | 2 | 6 | yes |
| UWS | M | 34 | 2 | 7814 | 1 | 0 | 1 | 1 | 0 | 2 | 5 | no |
| UWS | F | 49 | 2 | 277 | 1 | 0 | 2 | 2 | 0 | 1 | 5 | yes |

